# Genomic characterization of the *C. tuberculostearicum* species complex, a ubiquitous member of the human skin microbiome

**DOI:** 10.1101/2023.06.16.545375

**Authors:** Nashwa M. Ahmed, Payal Joglekar, Clayton Deming, NISC Comparative Sequencing Program, Katherine P. Lemon, Heidi H. Kong, Julia A. Segre, Sean Conlan

**Affiliations:** Microbial Genomics Section, Translational and Functional Genomics Branch, NHGRI, NIH, Bethesda, Maryland, USA; NIH Intramural Sequencing Center, NHGRI, NIH, Rockville, Maryland, USA; Alkek Center for Metagenomics & Microbiome Research, Department of Molecular Virology & Microbiology, Baylor College of Medicine, Houston, Texas, USA; Division of Infectious Diseases, Texas Children’s Hospital, Department of Pediatrics, Baylor College of Medicine, Houston, Texas, USA; Cutaneous Microbiome and Inflammation Section, NIAMS, NIH, Bethesda, Maryland, USA

## Abstract

*Corynebacterium* is a predominant genus in the skin microbiome, yet its genetic diversity on skin is incompletely characterized and lacks a comprehensive set of reference genomes. Our work aims to investigate the distribution of *Corynebacterium* species on the skin, as well as to expand the existing genome reference catalog to enable more complete characterization of skin metagenomes. We used V1-V3 16S rRNA gene sequencing data from 14 body sites of 23 healthy volunteers to characterize *Corynebacterium* diversity and distribution across healthy human skin. *Corynebacterium tuberculostearicum* is the predominant species found on human skin and we identified two distinct *C. tuberculostearicum* ribotypes (A & B) that can be distinguished by variation in the 16S rRNA V1-V3 sequence. One is distributed across all body sites and the other found primarily on the feet. We performed whole genome sequencing of 40 *C. tuberculostearicum* isolates cultured from the skin of five healthy individuals across seven skin sites. We generated five closed genomes of diverse *C. tuberculostearicum* which revealed that *C. tuberculostearicum* isolates are largely syntenic and carry a diversity of methylation patterns, plasmids and CRISPR/Cas systems. The pangenome of *C. tuberculostearicum* is open with a core genome size of 1806 genes and a pangenome size of 5451 total genes. This expanded pangenome enabled the mapping of 24% more *C. tuberculostearicum* reads from shotgun metagenomic datasets derived from skin body sites. Finally, while the genomes from this study all fall within a *C. tuberculostearicum* species complex, the ribotype B isolates may constitute a new species.

**IMPORTANCE:** Amplicon sequencing data combined with isolate whole genome sequencing has expanded our understanding of *Corynebacterium* on the skin. Human skin is characterized by a diverse collection of *Corynebacterium* species but *C. tuberculostearicum* predominates many sites. Our work supports the emerging idea that *C. tuberculostearicum* is a species complex encompassing several distinct species. We produced a collection of genomes that help define this complex including a potentially new species which we are calling *C. hallux* based on a preference for sites on the feet, whole-genome average nucleotide identity, pangenomics and growth in skin-like media. This isolate collection and high-quality genome resource sets the stage for developing engineered strains for both basic and translational clinical studies.

Microbiomes are shaped by taxa that are both characteristic to those sites and functionally important to that community. The genus *Corynebacterium* is one such taxa for the human skin and nares. Foundational studies using 16S rRNA gene sequencing and shotgun metagenomics by our lab (1, 2) and others (3) have established *Corynebacterium* as common members of the skin microbiome. While *Corynebacterium* have been positively correlated with the resolution of dysbiosis associated with eczema flares (4), the importance of the Corynebacterium spp. is less defined for skin disease severity in primary immune deficient patients (5, 6). *Corynebacterium* spp. are predominant members of the human aerodigestive tract microbiome (nares, oral cavity and respiratory tract) (3) and participate in microbe-microbe interactions with members of nasal microbiome (7, 8). *Corynebacterium* have been shown to engage with the host immune system, specifically *C. accolens*-promoted IL23-dependent inflammation in mice on a high-fat diet (9). *C. bovis* and *C. mastiditis* have been shown to predominate the microbiome of a ADAM10-deficient mouse model (10) as well as an ADAM17-deficient mouse model of eczema (11). Finally, *C. tuberculostearicum* has been shown to induce inflammation in human epidermal keratinocyte cell cultures (12). These studies establish *Corynebacterium* spp. as key members of the skin microbiome capable of both microbe-microbe and microbe-host interactions.

A critical resource for understanding the biology of *Corynebacterium* on the skin is a robust collection of complete reference genomes, including isolates collected from a variety of individuals and body sites. Previously published genome collections from skin- or nares-resident species include *Staphylococcus epidermidis* (13), *Cutibacterium acnes* (14) and the recent comparative analysis of *Dolosigranulum pigrum* (15). Of note, while emerging bioinformatic methods and pipelines are now being employed to extract nearly-complete genomes (MAGs) from metagenomic assemblies of skin samples (16), MAGs are not yet a substitute for genomes from cultured isolates to understand strain level or pangenomic diversity. In addition to functional prediction, comparative genomics is increasingly being used to augment conventional microbiological methods to define or redefine taxonomic boundaries (17, 18), as well as describe the full extent of diversity within these boundaries (19). A pangenome, which encompasses the complete set of genes present within a set of genome sequences, enables the characterization of gene-level heterogeneity within a taxonomic group. The pangenome is commonly subdivided into the ‘core’ genome, referring to genes present in all strains, and the ‘accessory’ or ‘dispensable’ genome, referring to those present in only one or some isolates. (The accessory pangenome can be further subdivided to reflect a wider range of gene uniqueness, *e.g.* singletons.) Thorough characterization of taxa is limited by the availability of representative and high-quality genome assemblies. Unfortunately, with the exceptions of clinically relevant *Corynebacterium* spp. (*e.g.*, *C. diphtheriae*, *C. striatum* and *C. pseudotuberculosis*), the genus is inadequately sequenced, with 75% of species having fewer than six genomes. This includes common skin-associated species like *C. tuberculostearicum* with just five unique isolate genomes, only two of which are from skin.

This work seeks first to characterize the distribution of *Corynebacterium* across 14 skin sites from 23 healthy volunteers. The second goal of this work focuses on what we identify as the predominant skin *Corynebacterium* species, *C. tuberculostearicum*. We have sequenced 23 distinct *C. tuberculostearicum* strains (n=40 genomes before dereplication), a five-fold increase in the number of publicly available, unique genomes (n=5). In addition to short-read assemblies, we generated five complete genomes which, along with the type strain (DSM44922), demonstrate that *C. tuberculostearicum* genomes are largely syntenic and carry a number of methylation systems as well as a CRISPR/Cas system. Genes from the *C. tuberculostearicum* genomes in our collection fall into 5451 gene clusters comprising the species pangenome. This expanded pangenome, as compared to existing public references, improved the mapping of *C. tuberculostearicum* metagenomic reads from unrelated healthy volunteers. In addition, we have identified a distinct *C. tuberculostearicum* clade that is highly enriched on the feet that may represent a new species, tentatively designated *Corynebacterium hallux*.

## Results

### *Corynebacterium* spp. are predominant members of the healthy skin microbiome

To explore the tropism of *Corynebacterium*, we surveyed the microbial diversity of healthy human skin using existing 16S rRNA V1-V3 amplicon sequencing data (5, 20). Clinical samples were obtained from 23 healthy volunteers across 14 body sites: sebaceous (back, Ba; occiput, Oc; external auditory canal, Ea; retroauricular crease, Ra; manubrium, Mb; glabella, Gb), moist (inguinal crease, Ic; antecubital crease, Ac), dry (hypothenar palm, Hp; volar forearm, Vf), foot (toe nail, Tn; toe web, Tw; plantar heel, Ph) and (N)ares. An average of 10,000 sequences per sample were generated which yielded a total of 8334 amplicon sequence variants (ASV), or unique 16S rRNA gene signatures. After rarefying the dataset to an even depth, 5967 ASVs remained. As expected, the dominant genera identified on the skin, present in 94% of skin samples, were *Cutibacterium* (41% of reads, ASV1 is *C. acnes*)*, Staphylococcus* (9% of reads, ASV2 is *S. epidermidis*), and *Corynebacterium* (9% of reads, ASV3 is *C. tuberculostearicum*). The genus Corynebacterium was present in 96% of the skin sites sequenced, averaging 17% of reads. With a preference for moist over sebaceous skin sites (Fig. S1), *Corynebacterium* thrives in the humid, temperate environments of the feet and nares. While variation in species composition was observed between individuals, some sites and habitats displayed species enrichment at specific locations across multiple individuals (Fig. 1A). We observed that *C. accolens* was enriched in the nares, with a prevalence of 83-87% across nares samples and constituted an average of 33-41% of Corynebacterium reads. *C. afermentans* were enriched across feet sites, where they were present in 54% of samples and comprised an average of 17% of *Corynebacterium* reads. Most notably, however, we found that *C. tuberculostearicum* was present in 94% of body sites and was often the most abundant *Corynebacterium*. *C. tuberculostearicum* reads represented 67% of corynebacterial reads in the feet, 47% in dry sites, 58% in sebaceous sites, and 46% in the nares.

**FIG 1.**
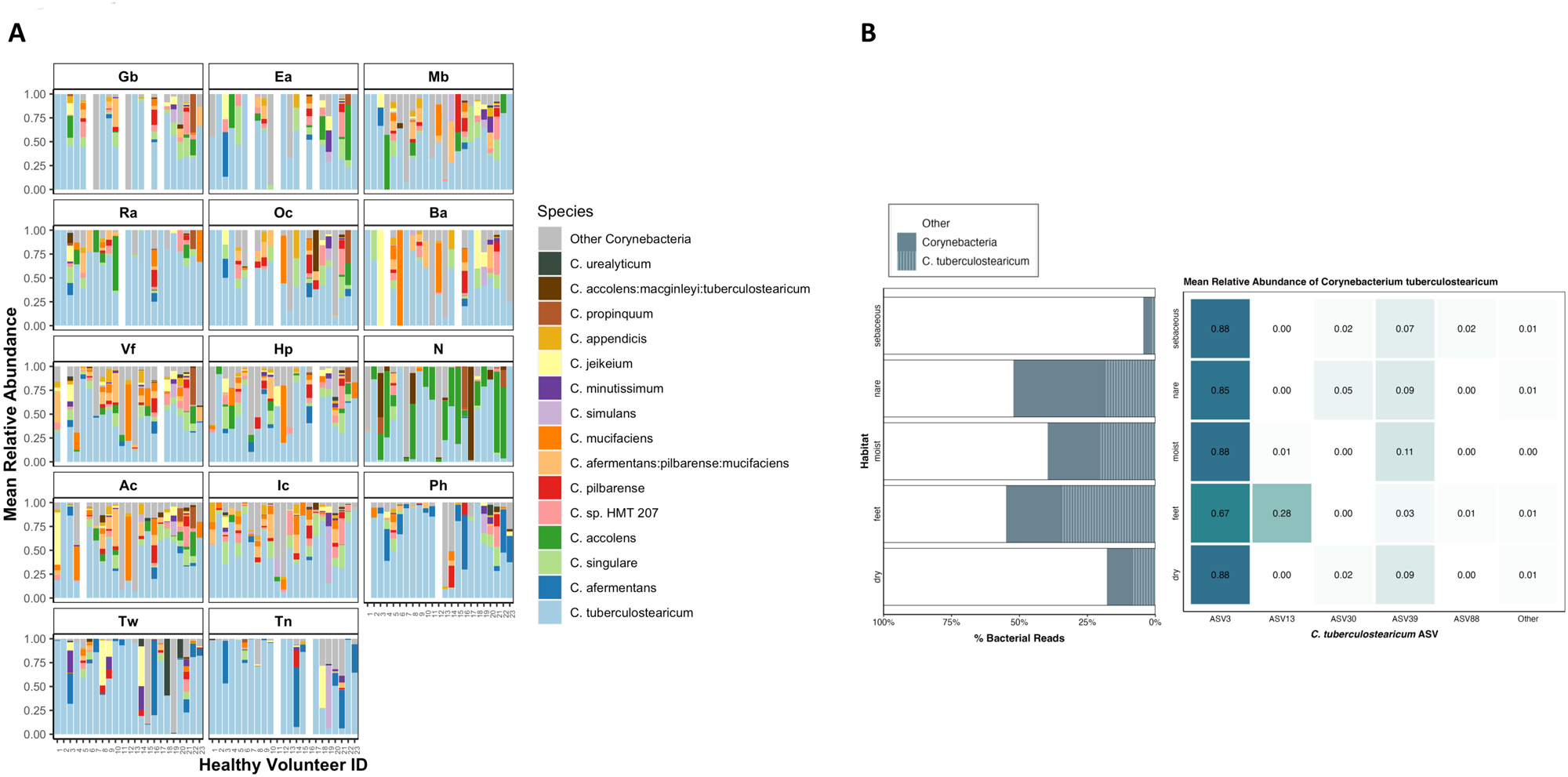
Corynebacterium species relative abundance in normal human skin microbiome. (A) Relative abundance of the 15 major Corynebacterium species across 14 skin sites: sebaceous (back, Ba; occiput, Oc; external auditory canal, Ea; retroauricular crease, Ra; manubrium, Mb; glabella, Gb), moist (inguinal crease, Ic; antecubital crease, Ac), dry (hypothenar palm, Hp; volar forearm, Vf), foot (toe nail, Tn; toe web, Tw; plantar heel, Ph) and (N)ares. Relative abundances determined by sequencing of the V1-V3 region of the 16S rRNA gene and subsetting to *Corynebacterium* reads. (B) Percent of total bacterial reads attributed to *Corynebacterium* and *C. tuberculostearicum* in each skin habitat. Of the six ASVs assigned to *C. tuberculostearicum*, mean relative abundance across skin habitats.

### *C. tuberculostearicum* is the most common skin Corynebacterium

A variety of marker gene approaches have been employed to determine the phylogenetic relationships between *Corynebacterium* species including combinations of 16S rRNA, *rpoB*, *rpoC* and *gyrA* genes (for review see (21)). In general, it is difficult to accurately classify *Corynebacterium* to the species-level using amplicon data and standard reference databases. The Human Oral Microbiome Database (3) is a curated database that includes a training set with a supraspecies taxonomic level enabling assignment of sequences to multiple species where ambiguity exists. In our case, >99.5% of sequences classified as *C. tuberculostearicum* using the Refseq classification, were also classified as *C. tuberculostearicum* (part of the *accolens*/*macginleyi*/*tuberculostearicum* superspecies) by eHOMD, including the two predominant 16S rRNA sequence variants, ASV3 and ASV13 which differed by a SNP and a single-base indel.

ASV3 constituted 83% of *C. tuberculostearicum*-classified reads (compared to < 8% for all other ASVs of this species) and showed a cosmopolitan distribution across body sites (Fig. 1B). Found in 100% of healthy volunteers (N=23) and 87% (254/293) of skin samples, ASV3 was predominant and ubiquitous across human skin. Relative abundance analysis revealed ASV3 abundance > 85% within all habitats except foot sites, where it made up 66% of *C. tuberculostearicum*-classified reads. As of this writing, the existing *C. tuberculostearicum* NCBI reference genomes containing complete V1-V3 sequences are all ASV3 as are 100% of 16S rRNA gene *C. tuberculostearicum* references in the SILVA reference database.

In contrast to cosmopolitan ASV3, ASV13 was enriched primarily on feet, constituting 28% of *C. tuberculostearicum*-classified reads from the Ph, Tw, and Tn sites (8%, 9%, and 70%, respectively). In 9 of 23 HVs, ASV13 constituted over 90% of *C. tuberculostearicum*-mapped reads within a single foot site (Fig S2); notably, much of this predominance was observed in Tn sites. In addition, we observed that some individuals exhibited within-site predominance by other less common ASVs, with some individuals colonized by a single non-dominant ASV across multiple body sites. In HV 12, for example, 52-100% of *C. tuberculostearicum*-mapped reads in each body site excluding the toenail are classified as ASV39. We also noted that, while sites on the feet (Ph, Tn, Tw) were often colonized by multiple ASVs, other body sites tended to be colonized by a single ASV.

We searched the SILVA database (v138.1) for perfect matches to the ASV13 sequence and found 152 matches, all associated with uncultured *Corynebacterium*. The majority of them were from our own full-length 16S rRNA gene sequencing of skin microbiome samples (1). This observation, combined with the fact that all the existing *C. tuberculostearicum* reference genomes had the more common ASV3 sequence variant, led us to hypothesize that the ASV13 sequence, which we hereafter refer to as ribotype B, could be associated with an unrecognized species or subspecies. For the purposes of the current work, we will use the term *C. tuberculostearicum* species complex (22, 23) to refer to all *C. tuberculostearicum*-like isolates found on skin. Additionally, we will refer to the predominant ASV3 OTU as ribotype A.

### Expanding the *C. tuberculostearicum* complex reference catalog

Prior to this study, only five *C. tuberculostearicum* species complex isolates had been sequenced and submitted to NCBI. Only two of those were from human skin and neither was a closed genome. To expand the *C. tuberculostearicum* complex reference genome catalog, we sequenced isolates from five different HVs (Supplementary Table S1). To enrich for the previously unsequenced ribotype B *Corynebacterium*, we screened skin-associated isolates by sequencing their 16S rRNA gene. This screen identified eight ribotype B isolates for further study. In total, we shotgun sequenced 40 isolates in the *C. tuberculostearicum* complex– 30 from ribotype A, 8 from ribotype B and 2 from other ASVs. Initial genome clustering using mash indicated that some of the isolates we sequenced were closely related. Therefore, we used dRep (24) to identify groups of highly similar genomes (ANImf > 99.5%) and chose the best representative genome for each genome set based on sequence assembly statistics: maximal N50, minimal number of contigs, and maximal genome size. In cases of comparable assembly quality, genomes were selected to increase body site representation. This resulted in a final set of 23 dereplicated *C. tuberculostearicum* complex genomes, with 18 from ribotype A, four from ribotype B, and one from ASV30.

### Whole-genome features of five complete *C. tuberculostearicum* complex genomes

In addition to a paucity of *C. tuberculostearicum* reference genomes at the time of this study, the ones that did exist were not associated with publications describing their general features. To address this, we selected five of the dereplicated genomes, three ribotype A and two ribotype B, for long-read sequencing on the PacBio platform. The subsequent finished or complete *C. tuberculostearicum* complex genomes revealed four copies of the 16S rRNA gene in each genome. For each genome, we performed a multi-sequence alignment containing each V1-V3 region copy along with the predominant ribotype A sequence first identified in our amplicon sequencing dataset. Within a genome, copies of the V1-V3 region are almost entirely identical across alignment, with the ribotype A *C. tuberculostearicum* complex genomes carrying four copies of ASV3. Admittedly, one exception was a single nucleotide variant identified in one copy of 16S rRNA 5’ region of CTNIH12 (Fig. S3). Notably, no variation was found in the ribotype B genomes, which both carried four identical gene copies marked by the two characteristic sequence variants as identified in the amplicon sequencing dataset. The within-genome homogeneity of 16S rRNA genes confirmed its usefulness as a marker.

These complete *C. tuberculostearicum* complex genomes also enabled us to directly compare the type strain (DSM 44922/FDAARGOS_1117; human bone marrow) to our ribotype A and B isolates without the ambiguity introduced by unfinished genomes. Supplemental Figure 4 shows that the five PacBio genomes from this study were largely co-linear, with >80% of the genome in large syntenic blocks, with the type strain DSM 44922. All five of the genomes in this study had a 440 kb region that was reorganized relative to the type strain. This region, comprising around 17.6% of the genome encoded 392 genes (387 coding). The breakpoints for inversions or translocated blocks in the reference were marked by mobile element families (*e.g.*, IS3, IS256, IS481, IS6) that suggest a mechanism for these rearrangements.

We extracted the methylation profiles from the PacBio reads of our five genomes (Supplementary Table S2). The most common methylation pattern, found in all five genomes, was N6-methyladenine modification (m6A) of GATC motifs (∼16,000 sites/genome; 99% methylated), typically associated the Dam methylase. A second motif AAAAC was also found to be methylated (m6A) in all five genomes (∼75% methylated). In addition to these two ubiquitous methylation patterns, ribotype A isolate CTNIH10 had two additional methylated motifs present in hundreds of copies (GGCANNNNNATC, GATDNNNNTGCC). CTNIH20 had an additional three methylated motifs present at 520-1626 sites/genome. Finally, CTNIH23 had evidence of an additional five methylation motifs across the genome that were all >98% methylated and present at 225-2159 sites/genome. Methylation systems are important for horizontal gene transfer, phage resistance and potential recombinant engineering of these strains.

The presence of CRISPR-Cas as well as other phage defense systems pose additional barriers to horizontal gene transfer (HGT). We detected an eight-gene Type I-E Cas gene cluster and two large repeat arrays (24 and 19 spacers) in the CTNIH20 ribotype B genome, but not in any of the other full-length genomes from this study. Additional CRISPR-Cas systems were detected in the short-read assemblies of CTNIH9 (ribotype A; Type I-E Cas gene cluster, 8 spacer CRISPR) and CTNIH22 (ribotype B; Type I-E Cas gene cluster, 12 spacer CRISPR). Prior to this, the only public *C. tuberculostearicum* complex reference genome with a CRISPR-Cas system was strain SK141 (ACVP01). A variety of other defense systems including restriction modification systems were also identified using the DefenseFinder tool (Supplementary Table S2) (25, 26).

Plasmids are important for the mobilization of virulence factors, antibiotic resistance genes and as tools for recombinant engineering. A single 4.2 kb plasmid was deposited in the public databases associated with the *C. tuberculostearicum* species complex, p1B146 (NC_014912) (27). Across the five long-read genomes sequenced here, we detected five plasmids, none of which aligned to p1B146. Two plasmids, pCT3-020e and pCT4-9116 from CTNIH23 and CTNIH12, had the same backbone as the *C. diphtheriae* plasmid pNG2 (*ORF9-traA-ORF11-parAB-repA*) but lacked the erythromycin resistance cassette. CTNIH20 carries three plasmids ranging in size from 21.2 kb to 27.3 kb. All three carried a *traA*/*recD2* ortholog encoding a relaxase/helicase but are otherwise unrelated. While most of the proteins on these three plasmids were annotated as hypothetical, pCT1-afe7 carried an *ebrB* efflux pump and pCT1-0563 carried an *stp* (spectinomycin/tetracycline) efflux pump predicted to be involved in resistance to dyes and antibiotics. All five plasmids were characterized by the presence of a TraA/RecD2 encoding gene, suggesting a common mobility mechanism. Furthermore, we found fragments of these plasmids in many of the contig-level genomes. For instance, the *C. tuberculostearicum* CIP 102622 genome (JAEHFL01) carried both the *stp* gene and a nearby transcription factor (>99.6% identity) on a 9.8 kb contig, showing the value of these plasmid references for identifying HGT elements.

### Taxonomic structure of the skin-associated *C. tuberculostearicum* species complex

While 16S rRNA amplicon sequencing enabled us to group *C. tuberculostearicum* species complex isolates into two predominant ASVs, the dereplicated genomes enable further high-resolution taxonomic analysis of this species complex. We used GET_HOMOLOGUES to extract core genes and build a phylogenetic tree based on core genome SNPs (Fig. 2). We noted additional taxonomic structure particularly amongst ribotype A isolates. The five public reference genomes were in the ribotype A-dominated portion of the tree as expected based on their 16S rRNA gene sequence.

**FIG 2.**
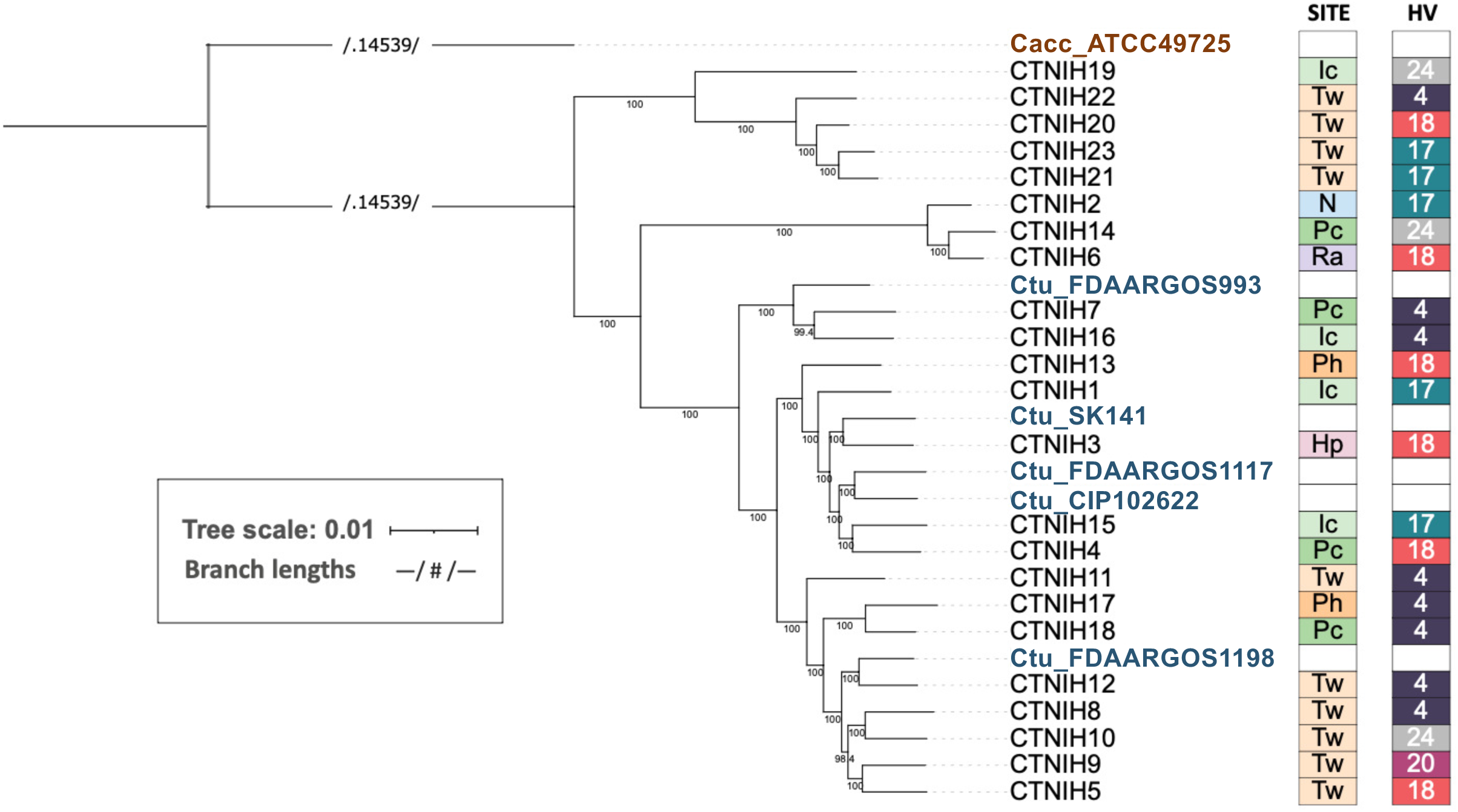
A maximum-likelihood phylogenetic tree of *C. tuberculostearicum* species complex genomes from this study and publicly available, calculated from 1315 core gene cluster alignments. Bootstrap values (located along internal nodes) were calculated from 1000 replicates. Clustering was generated using GET_HOMOLOGUES OrthoMCL v1.4 option with minimum coverage 90% in BLAST pairwise alignments. The tree was rooted on outgroup *C. accolens* ATCC 49725. On the right of tree, boxes depict site (body site locations defined in Figure 1) and individual (HV) from which each isolate was cultured. Sites are colored by niche type, with moist in shades of green; feet in shades of orange; dry in pink; sebaceous in lavender; and nares in blue. Individuals are randomly but consistently colored.

While there was good correlation between 16S rRNA ASVs and the core SNP tree, we noted a single isolate, CTNIH19, which carried a ribotype A allele but localized with the ribotype B isolates on the tree. CTNIH19 was isolated from the inguinal crease and is the most basal member of this clade. Work by Cappelli and colleagues recently defined a number of new Corynebacterium species and CTNIH19 was >99.9% identical to a species they designate *C. curieae* (28). We calculated the average nucleotide identity across isolates using pyANI (Fig. S5) and determined that ribotype B isolates share ANI >97% with themselves and <94% with other *C. tuberculostearicum* complex genomes, including *C. curieae*. We submitted our ribotype B isolate genomes to the DSMZ type strain genome server (TYGS) (29) to obtain a taxonomic/nomenclature assignment. TYGS predicted that ribotype B genomes belong to a new species in both the whole genome and 16S rDNA trees. The closest TYGS references were *C. tuberculostearicum* DSM 44922 and *C. kefirresidentii*. *C. keffiresidentii* was first described in 2017 (30) after isolation from kefir grains but has not been accepted as an official species yet. Three of our isolates (CTNIH2, CTNIH6, CTNIH14) from three different healthy volunteers were 98% identical to the *C. kefirresidentii* reference, calling into question whether kefir is the only natural host for this bacterium (22, 23).

### Pangenome of the skin-derived *C. tuberculostearicum* complex

We performed a pangenomic analysis to describe the coding diversity of the *C. tuberculostearicum* species complex (Figs. 3). We generated an anvi’o (31) pangenomic map to illustrate genomic variation across the combined set (N=28) of NCBI reference genomes and our dereplicated genomes. (Fig. 3A). Pangenome openness was estimated using the Heap’s law model (Fig. 3B) as proposed by Tettelin et al (32). The model indicated an open pangenome (0.30, ± 0.1, γ > 0), predicting that the *C. tuberculostearicum* pangenome would increase with more genomes analyzed. With our additional 23 genomes, the total pangenome size increases from 3080 genes to 5451 genes, resulting in an expansion of the non-core, or accessory genome by over 300% (Fig. 3C). We performed a functional characterization of 23 lab-sequenced and 5 NCBI-derived *C. tuberculostearicum* species complex genomes using the eggNOG-mapper annotation tool (Fig. S6), which returned annotations for 83.2% of orthologous gene clusters (of which 21% are annotated COG category “S”, *Function Unknown*). Interestingly, among the non-core genes, inorganic ion metabolism and transport-related genes were among the most abundant. We performed a principal components analysis (PCA) of gene presence/absence data describing our 23 genomes and 2 skin-derived reference sequences (Fig. 4A). We observed site-specific clustering of genomes isolated from the feet and moist environments. In addition, we observed distinct clustering of ribotype B isolates (circles) away from other foot-derived *C. tuberculostearicum* complex genomes, which agreed with the core phylogenetic clustering. In addition, we identified 11 genes (A=2, B=9) that were unique to and carried by every member of a ribotype (Supplementary Table. S3), four of which we were able to assign functional annotation using the UniprotKB sequence similarity search tool, including a bacteriocin and ferric uptake protein.

**FIG 3.**
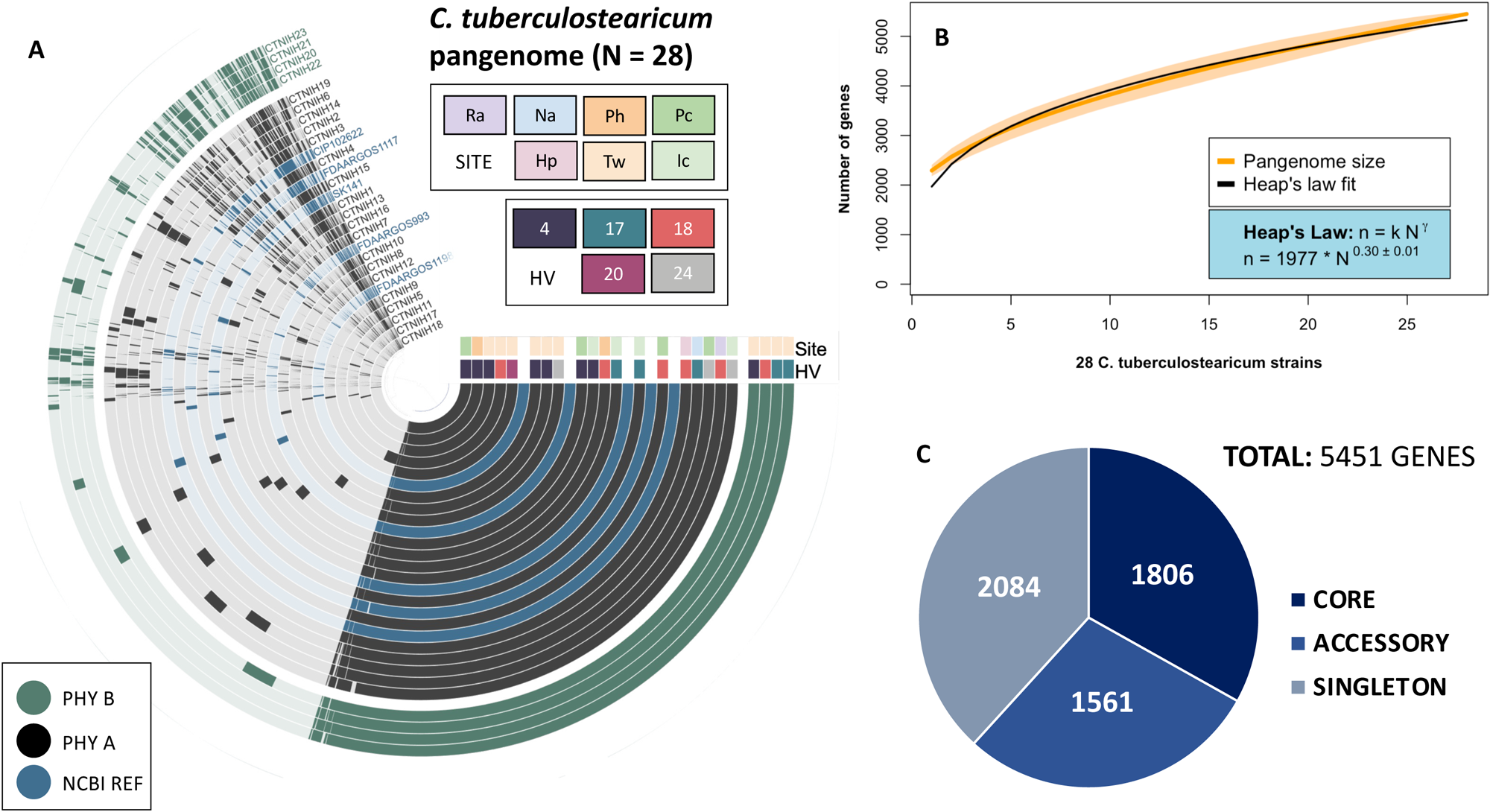
The *Corynebacterium tuberculostearicum* pangenome. (A) Anvi’o pangenomic map for 28 *C. tuberculostearicum* genomes (including 5 NCBI reference genomes). Genomic rings are annotated by skin site and HV (healthy volunteer) metadata and ordered by pyANI average nucleotide identity (ANIb). Genome margins are manually adjusted for clarity. (B) Heap’s Law estimate of pangenome openness for 28 genomes. A rarefaction curve showing the total number of genes accumulated with the addition of new genome sequences in random order with 1000 permutations. Shaded regions represent the 95% confidence interval. A Heap’s law model was fit to the resultant curve to calculate k and γ values (1977±38.0 and 0.30±0.01, respectively). (C) Number of core (belonging to all genomes), accessory (belonging to two or more genomes), and singleton (belonging to only one genome) genes. The expanded pangenome contains 5451 genes using 90% sequence identity as a cutoff parameter.

**FIG 4.**
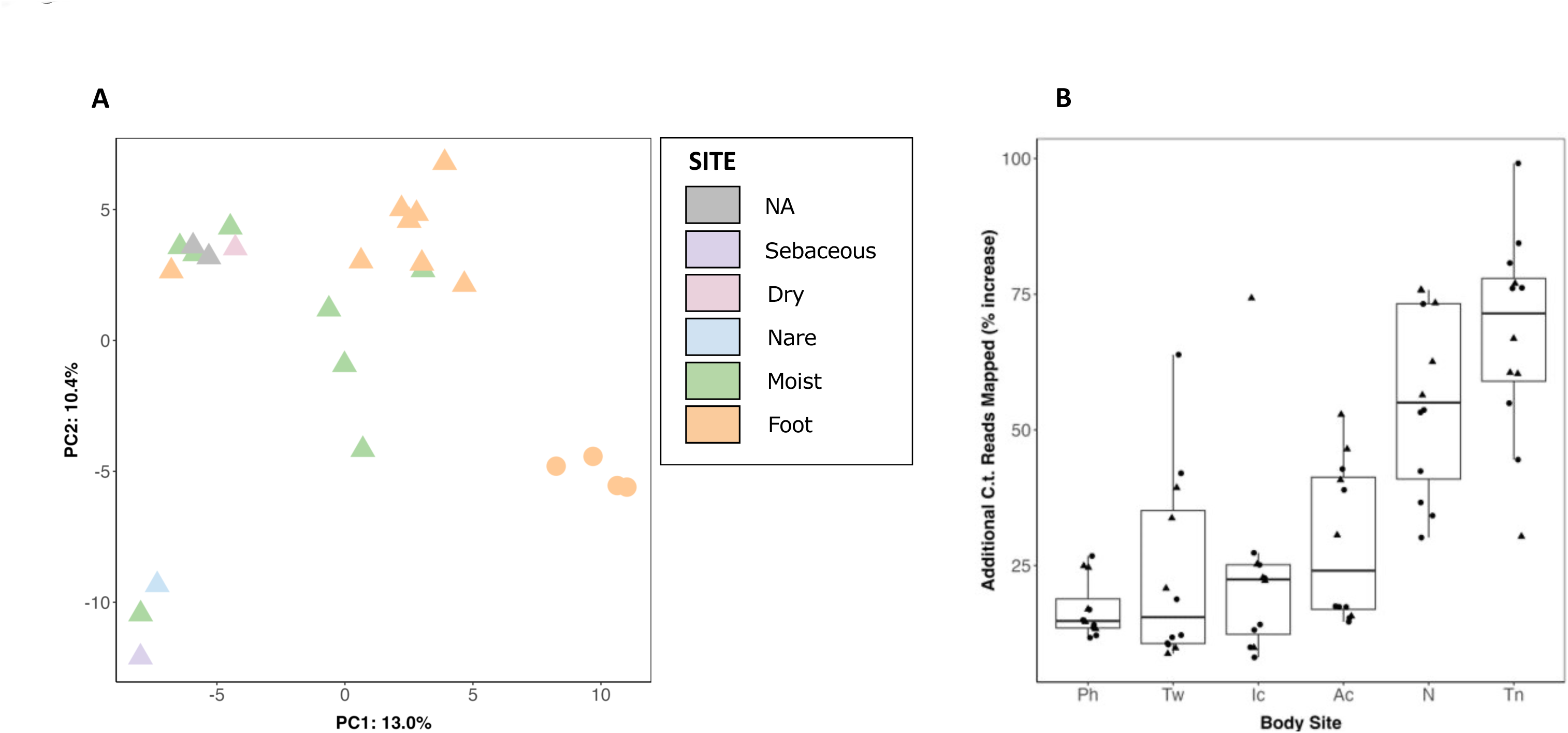
*C. tuberculostearicum* complex pangenome clustering and improved metagenomic read mapping. (A) Principal components analysis of orthologous gene clustering. The gene presence/absence data for 25 genomes (including two NCBI references, shown in gray) was analyzed using principal components analysis. Ribotype B genomes are shown as circles; other genomes are triangles. (B) Improvement in shotgun metagenomic read mapping with a 28 member *C. tuberculostearicum* database as compared to the 5 member NCBI database. Percent increase in mapped *C. tuberculostearicum* reads by body site. Each point is a healthy volunteer. Triangles mark healthy volunteers that contributed one or more isolates to the expanded mapping database.

### Improved metagenomic read mapping using an expanded *C. tuberculostearicum* pangenome

We tested whether the expanded genomic reference set could improve the rate of *C. tuberculostearicum* read mapping in a set of metagenomic datasets from 12 healthy volunteers at 6 body sites (2). Reads were mapped with bowtie2 (33) against a genome database consisting of the five unique NCBI references or a database of the NCBI references plus the dereplicated genomes from this study. Overall, 27% more reads were assigned to *C. tuberculostearicum* using the expanded genome set as compared to NCBI references alone. While the five HVs with isolate genomes in the expanded database showed slightly better classification, median improvement of 32%, over those without isolates in the database (median=24%), the difference was not statistically significant, showing the broad utility of these genomes. Furthermore, when broken down by body site, toenail (Tn) sites showed the largest improvement in *C. tuberculostearicum* read assignment (72%) while nares, which only contributed a single genome to the expanded database, improved by 55% (Fig 4B). To control for spurious read mapping to repetitive elements or other assembly artifacts, these analyses were repeated using only the predicted gene catalogs, rather than the whole genomes, and very similar improvements in *C. tuberculostearicum* read mapping were observed, 25% median improvement and a similar site-dependence.

### Growth of skin-derived *C. tuberculostearicum* in sweat media

Members of the *C. tuberculostearicum* species complex are widely distributed across the skin’s microenvironments. Differences in body site physiology and nutrient composition inherent to each niche may provide selective growth advantages (and disadvantages) to a subset of strains. We performed a pilot experiment to investigate differential growth phenotypes of *C. tuberculostearicum* species complex ribotypes in skin-like media. (Fig 5) *Corynebacterium* are often cultured on brain-heart infusion (BHI) media plates supplemented with 1% Tween-80 (BHI + 1% Tween80) so this media was used as a positive control for growth in liquid medium (Fig. 5B). Isolates were cultured on two types of medium consisting of a complex mixture of amino acids, lipids, and other metabolites that mimic human eccrine sweat, with one medium supplemented to include a sebum-like synthetic lipid mixture. Both simulated sweat medias were supplemented with 0.1% Tween-80. A collection of eight skin-derived *Corynebacterium* strains consisting of four ribotype A and four ribotype B strains were grown for 20 hours in triplicate and in two separate experiments for each strain and medium condition (N=6) (Fig. 5 B-D). In all three growth conditions, ribotype B isolates demonstrated a lower mean OD_600_ over time than ribotype A isolates (Fig. 5A). Using ANOVA and the Tukey method, we determined that the area-under-the-curve (AUC) difference between the two ribotypes is statistically significant (p <0.0001) for all media conditions. This pattern was particularly pronounced in the BHI + 1% Tween80 and Sweat media + 0.1% Tween80 conditions. We observed that the addition of synthetic lipid mixture to the eccrine sweat-like medium attenuated, however still maintained the growth difference between ribotype B and other strains, suggesting lipid-limited growth for members of ribotype B.

**FIG 5.**
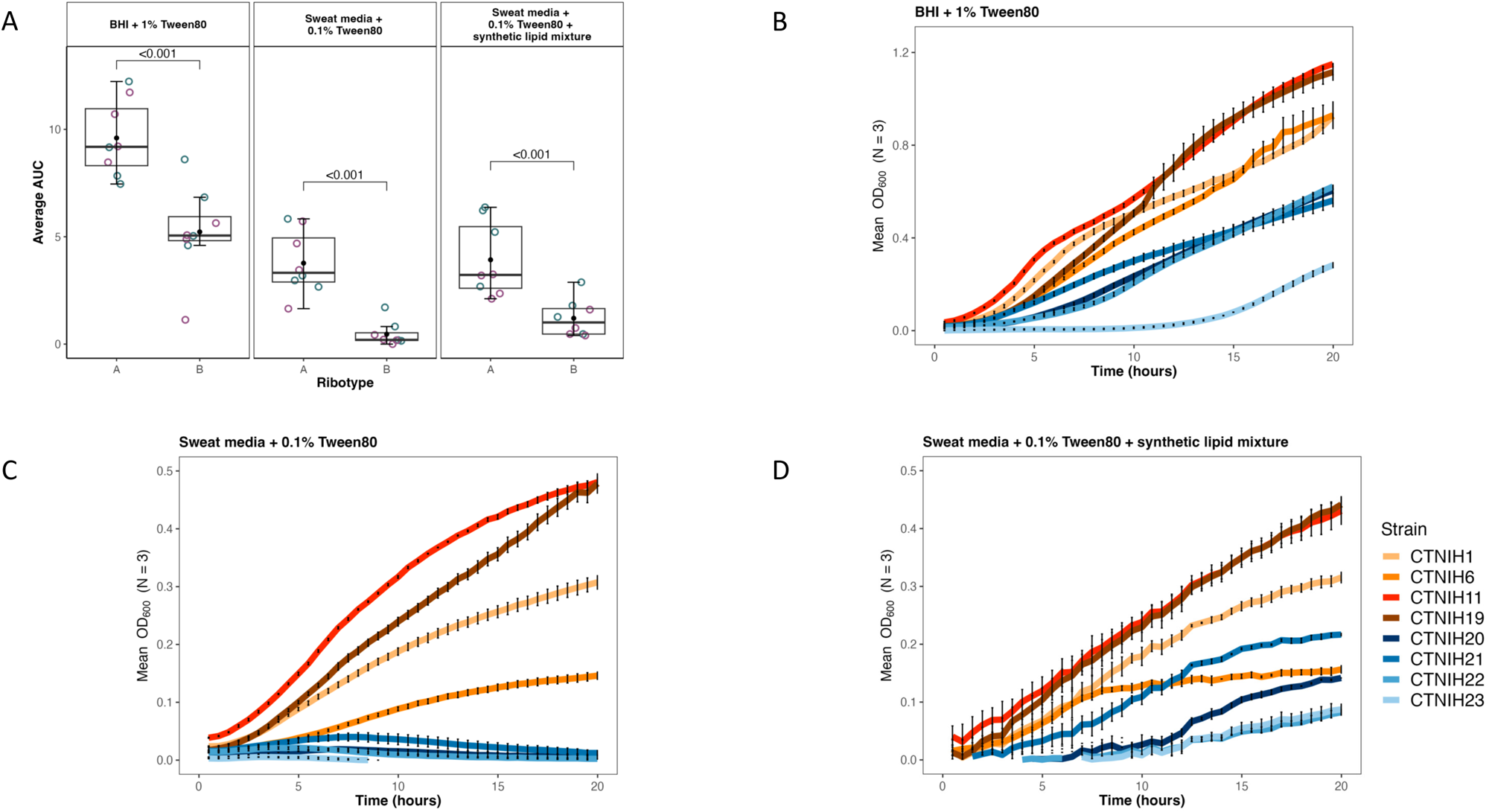
Growth phenotypes of select *C. tuberculostearicum* complex strains in synthetic sweat media. (A) Empirical area under curve comparison of *C. tuberculostearicum* species complex strains from ribotype A and ribotype B, with biological replicates grouped by color. Strains were grown in Brain Heart Infusion (BHI) + 1% Tween; Sweat media + 0.1% Tween80; Sweat media + 0.1% Tween80 + synthetic lipid mixture. Medium composition is described in further detail in Methods. (B-D) Selected growth curves from a representative experiment plotted with standard error. Ribotype B isolates are shown in shades of blue; Ribotype A isolates are shown in shades of red.

## DISCUSSION

In this study, we investigated the genomic diversity of the predominant yet under-sequenced Corynebacterium genus. Our survey of microbial diversity across human skin revealed niche-specific enrichment of Corynebacterium species and identified *C. tuberculostearicum* as a predominant and widespread species on human skin. Our amplicon-based analysis was able to identify a site-specific novel 16S rRNA gene ribotype which led to an expanded sequencing of the *C. tuberculostearicum* species complex. In total, we sequenced 23 distinct isolates belonging to the *C. tuberculostearicum* species complex including *C. tuberculostearicum* (n=15), *C. kefirresidentii* (n=3), *C. curieae* (n=1) and a novel species we are calling *C. hallux* (n=4). Discovery of *C. kefirresidentii* on human skin and nares suggests that humans are a natural host for this species.

*C. hallux* is likely a new species of skin-associated *Corynebacterium* and merits further work to formally name it. It was cultured from three different healthy volunteers, detected by amplicon sequencing in most HVs, represented in the recently published SMGC (SMGC_122) (16) and detected in public 16S rRNA gene databases entries associated with skin. In our healthy volunteers, it was enriched in sites on the feet, particularly the toenail and toe web. Microbial communities on the feet are highly diverse and relatively unstable (2) subject to temperature fluctuations and invasion by environmental microorganisms.

This study helps to resolve the diversity of *C. tuberculostearicum* species complex strains and provides an important genetic resource for future study. Our whole-genome sequencing uncovered insights into the genetic diversity of the complex and improved read-mapping overall by >24%, which will in turn bolster future sequencing efforts and lead to better characterization of *Corynebacterium* across human skin. While our bioinformatic analysis greatly expands the non-core genome, a significant proportion of these genes are putative and lack definitive annotation. Overall, we did not detect obvious gene-level differences between ribotype B and other strains that would explain the observed differences in site distribution pattern and growth on synthetic media. Only 11 genes perfectly segregated the two ribotypes and limitations of functional annotation tools resulted in only hypothetical functional annotations.

Our pangenomic analysis did not reveal major metabolic pathways or modules that differed between ribotype A and B isolates that would explain niche specificity, however there were two examples of genes with the potential to affect within-niche competition. One of the genes specific to ribotype B shared sequence similarity with a Lactococcin 972 family bacteriocin. Bactericidal activity of ribotype B against closely related strains could contribute to patterns of within-site dominance as observed between ribotypes (Fig. S2). Bactericidal peptides have recently gained interest as a possible therapeutic intervention for gastrointestinal disease (34). Furthermore, *Corynebacterium* have been shown to be enriched in a recent study (35) of post-operative, healing wounds, suggesting an opportunity for biotherapeutic applications. We also identified a ribotype B-unique copy of a gene encoding ferrous iron transport protein B, a major regulator of bacterial iron uptake. Iron is an essential nutrient for survival, requiring the development of highly-efficient sequestering mechanisms by pathogenic and avirulent bacteria alike (36, 37). Under conditions of limited nutrient bioavailability, enhanced ferric uptake may prove to be a determining factor of intraspecies competition.

On both rich and skin-like media, we observed that ribotype B strains grew less robustly compared to other strains. Thus, different strains may perform unique roles within their respective niches. The observed strain-specific distribution pattern may arise from selective growth advantages including differences in nutritional requirements or nutrient acquisition mechanisms between strains. Understanding the mechanisms of this variability has important clinical implications. For example, further characterizing the nutritional limits for sustained growth may lead to prebiotic therapeutics to augment the growth of beneficial strains within a given microenvironment, or engineering site-specific, microbe-based drug delivery systems. Understanding the roles and requirements of host-associated microbial communities in maintaining skin health will provide insight into the emergence of skin disorders in addition to novel therapeutic interventions to combat them.

## Materials and Methods

### Subject recruitment and sampling

Healthy adult male and female volunteers (HVs) 18–40 years of age were recruited from the Washington, DC metropolitan region. This natural history study was approved by the Institutional Review Board of the National Human Genome Research Institute (clinicaltrials.gov/NCT00605878) and the National Institute of Arthritis and Musculoskeletal and Skin Diseases (https://clinicaltrials.gov/ct2/show/NCT02471352) and all subjects provided written informed consent prior to participation. Sampling was performed as described previously (20).

### 16S rRNA gene sequencing

16S rRNA gene amplicon sequencing of these samples has been described previously (5). Briefly, each DNA sample was amplified with universal primers flanking variable regions V1 (27F, 5′-AGAGTTTGATCCTGGCTCAG) and V3 (534R, 5′-ATTACCGCGGCTGCTGG). For each sample, the universal primers were tagged with unique indexes to allow for multiplexing/demultiplexing (38). The following PCR conditions were used: 2 μl 10X AccuPrime Buffer II, 0.15 μl Accuprime Taq (Invitrogen, Carlsbad, CA), 0.04 μl adapter+V1_27F (100 μM), 2 μl primer V3_354R+barcode (2 μM), and 2 μl of isolated microbial genomic DNA. PCR was performed in duplicate for 30 cycles followed by PCR-clean up and amplicon pooling of ∼10 ng DNA. Duplicate amplicons were combined, purified (Agencourt AMPure XP-PCR Purification Kit (Beckman Coulter, Inc., Brea, CA)), and quantified (QuantIT dsDNA High-Sensitivity Assay Kit (Invitrogen, Carlsbad, CA)). An average of ∼8 ng DNA of 94 amplicons were pooled together, purified (MinElute PCR Purification Kit (Qiagen, Valencia, CA)) and sequenced on a Roche 454 GS20/FLX platform with Titanium chemistry (Roche, Branford, Connecticut). Flow-grams were processed with the 454 Basecalling pipeline (v2.5.3).

### 16S rRNA gene amplicon analysis

Sequencing data were processed using DADA2 v1.20.0 (39). Sequences were filtered and trimmed as recommended by the software developers and truncated to 375 nt: filterAndTrim(fnFs, filtFs,maxN=0, maxEE=c(2), truncQ=2,truncLen=c(375)). Sample inference was performed using the learnErrors (randomize=TRUE) and the dada (HOMOPOLYMER_GAP_PENALTY=-1, BAND_SIZE=32) commands. Chimeras were removed using removeBimeraDenovo (method=“consensus”, allowOneOff=TRUE). Taxonomy was assigned using assignTaxonomy (minBoot=70) command in DADA2 with the Refseq (https://zenodo.org/record/3266798) or eHOMD v15.1 V1V3 (3) training set databases. The resulting amplicon sequence variants (ASVs), taxonomy and sample metadata were used to build a phyloseq (40) object that was used for further analysis.

### Bacterial culturing

*Corynebacterium* isolates were cultured from healthy volunteers as previously described (16). Briefly, skin samples were collected with eSWabs (COPAN e480C) in liquid Amies. Samples were diluted and plated on brain heart infusion agar with 1% Tween 80. Potential *Corynebacterium* isolates were taxonomically classified by amplifying and Sanger sequencing the full length 16S rRNA gene with primers (8F, 5′-AGAGTTTGATCCTGGCTCAG) and (1391R, 5′-GACGGGCGGTGWGTRCA).

### Bacterial whole genome sequencing

Genomic DNA was purified for each isolate, from which Nextera XT (Illumina) libraries were generated. Each isolate was sequenced using a 2×151 paired-end dual index run on an Illumina NovaSeq 6000. The reads were subsampled to achieve 80-100x coverage using seqtk (version 1.2), assembled with SPAdes (version 3.14.1) (41) and polished using bowtie2 (version 2.2.6) and Pilon (version 1.23) (42). To achieve full reference genomes for select isolates, genomic DNA was sequenced on the PacBio Sequel II platform (version 8M SMRTCells, Sequel II version 2.0 sequencing reagents, 15 hr movie collection). The subreads were assembled using Canu v2.1 and polished using the pb_resequencing workflow within PacBio SMRTLink v.9.0.0.92188. Genome annotation was performed using National Center for Biotechnology Information (NCBI) Prokaryotic Genome Annotation Pipeline (PGAP: https://www.ncbi.nlm.nih.gov/genome/annotation_prok/). Methylation patterns for the assembled genomes were determined using the pb_basemods workflow in SMRTLink v.9.0.0.92188. Whole genome and plasmid alignments were generated in mummer (v3.9.4alpha) and visualized in R.

Full-length 16S rRNA gene copies were extracted from each PacBio complete genome. Briefly, reference *Corynebacterium* 16S rRNA sequences were downloaded from the RDP database (Good quality, >1200 nt) and used as a BLAST database to identify the coordinates of the four copies in the genome. To detect intragenomic variation in the 16S rRNA gene, all copies within each genome were compared against each other using the EMBL-EBI Multiple Sequence Alignment Tool (MUSCLE). Whole genome alignments were generated in Mauve v 2.4.0.

### Phylogenetic analysis

Publicly available genomes were downloaded from NCBI including *C. tuberculostearicum* (CP068156, CP06979, CP065972, ACVP01, JAEHFL01), *C. kefirresidentii* (CP067012, JAHXPF01), *C. curieae* (JAKMUU01) and *C. accolens* (ACGD01). GET_HOMOLOGUES (v09212021) was used to cluster protein sequences from 29 genomes (28 *C. tuberculostearicum*, 1 *C. accolens*) into orthologous groups and generate a core gene alignment. Prokka GBK files were used as input for clustering. The OrthoMCL (v1.4) option was used to group sequences utilizing the Markov Clustering Algorithm with a minimum coverage value of 90% in blast pairwise protein alignments. A strict core consensus genome was generated by calculating the intersection of single copy genes present in all 29 genomes. The accompanying GET_PHYLOMARKERS (v. 2.2.9.1) pipeline was used to identify markers for phylogenetic inference. IQTREE (v 2.1.2) was used to generate a maximum-likelihood phylogenetic tree from marker gene cluster alignments with 1000 bootstrap replicates. and a mean branch support value cutoff of 0.7. The top-scoring tree was visualized and annotated using the web-based program interactive Tree of Life (iTOL v6). The average nucleotide identity (ANIb) matrix for all sequences was plotted and annotated using the package heatmap.2/R.

### Gene calling and Annotation

The Prokka (v1.14.6) pipeline was used for gene calling and annotation. GFF3- and GBK-format annotations were generated for 28 *Corynebacterium tuberculostearicum* sequences derived from 23 lab isolates and 5 NCBI references (CIP 102622, FDAARGOS 993, FDAARGOS 1117, FDAARGOS 1198, SK141), in addition to the *Corynebacterium accolens* representative genome (ATCC 49725) sequence.

### Pangenome calculation

Three *C. tuberculostearicum* pangenomes (for all reference sequences, skin-derived reference sequences and lab sequences, and all sequences) were calculated from Prokka-derived GFF3 files using Panaroo on sensitive mode with a sequence identity threshold of 90% and otherwise default parameters. The resultant gene presence/absence tables were used for downstream analysis.

### Pangenome visualization

A pangenomic map was created using anvi’o (v. 7.1) with imported Prokka gene calling information and annotations (GBK format) for 23 lab-sequenced and 5 NCBI reference *C. tuberculostearicum* genomes. Strains were annotated with sample metadata including skin site, general skin habitat, healthy volunteer ID, as well as phylogenetic grouping from GET_HOMOLOGUES analysis. Average nucleotide identity (ANIb) of aligned regions was calculated within anvi’o using pyANI. In addition, eggNOGG (v. 2.1.7) gene annotations were used for gene cluster annotation with the NCBI COG Database (2020) and visualized using ggplot2/R. Core, accessory, and singleton gene counts were derived from gene presence/absence tables for (1) all reference sequences and (2) all sequences. Counts were visualized as pie charts.

A gene rarefaction curve for the *C. tuberculostearicum* pangenome was found by applying the Vegan/R specaccum() function to the gene presence/absence table, with a random order of additions of genomes permuted 1000 times. A Heap’s law power law model was fitted to the curve using the nls function in stats/R to calculate constants K and α. The curve was visualized using ggplot2/R.

A principal components analysis was performed on the gene presence/absence table using the prcomp function in stats(v 3.6.2)/R. The resultant object was visualized using ggplot2/R.

### Unique genes

Scoary (v. 1.6.16) was used to identify genes unique to ribotype A and B for 23 lab-sequenced *C. tuberculostearicum* complex isolates. Gene sequences were queried using the UniProtKB online sequence similarity BLAST tool (https://www.uniprot.org/blast).

### Metagenomic read mapping

Metagenomic reads from 12 healthy volunteers at 6 body sites, adapter trimmed and host subtracted as described in (2), were aligned to a bowtie2 (v 2-2.4.5) database built from five NCBI *C. tuberculostearicum genomes* (CIP_102622, DSM 44922, FDAARGOS_1198, FDAARGOS_993, SK141) with or with supplementation with 23 non-redundant genomes from this study; default bowtie2 parameters were used: --end-to-end -- sensitive.

### Growth curve starter cultures

Isolates for differential growth analysis were selected on the basis of i) reliable growth in Brain-Heart-Infusion+Tween80 and ii) coverage of the phylogenetic tree. *C. tuberculostearicum* isolates were grown in overnight liquid culture consisting of BHI broth (Sigma-Aldrich), augmented with 1% RPI Tween80, and 40ug/ml Fosfomycin (BHI-T-F) at 37°C with shaking at 220 rpm. To make the “Sweat media + 0.1% Tween80” media, we filter-sterilized Pickering Artificial Eccrine Perspiration Cat. No. 1700-0023 (pH 6.5) and RPI Tween80 (1%). This medium was then vortex-combined with 1% volume of synthetic apocrine sweat (Pickering Cat. No. 1700-070X) to produce the “Sweat media + 0.1% Tween80 + synthetic lipid mixture” medium.

### Differential growth experiments

*C. tuberculostearicum* liquid cultures were pelleted, washed and diluted 10-fold in diH2O to an OD_600_ of ∼0.1. Differential media were inoculated with diluted culture at a concentration of 100:1 and plated in triplicate across a 96-well microplate. Bacterial growth was recorded using the Epoch 2 Microplate. OD_600_ readings were taken at 30 minute intervals throughout a 24-hour time span. The experiment was performed in duplicate. OD_600_ measurements were exported, corrected via blank subtraction, and plotted using ggplot2/R. The package growthcurver/R was used to calculate empirical area under the curve (AUC) for all isolate:media combinations. Statistical significance testing for ribotype:media interactions were performed using ANOVA and a post-hoc Tukey test.

### Data Availability

Genome data are deposited under the NCBI BioProjects PRJNA854648, PRJNA694925 and PRJNA854648 (see Table S1). Some amplicon data were published previously (n=145; PRJNA46333)(5) and the remainder are new to this study (n=168; PRJNA46333)

## ACKNOWLEDGEMENTS

We thank Tommy Hiller Tran for valuable bioinformatics discussions. We thank Dr. Matthew Kelly for discussions of *Corynebacterium* biology. The computational resources of the NIH High-Performance Computation Biowulf Cluster (http://hpc.nih.gov) were used for this study. This work was supported by the Intramural Research Programs of the National Human Genome Reseach Institute and the National Institute of Arthritis and Musculoskeletal and Skin Diseases.

N.A. performed bioinformatics and growth curve analyses, prepared figures and wrote the manuscript; P.J. provided technical support for growth curve experiments; C.D. cultured bacteria and prepared DNA sequencing libraries; NISC sequenced amplicon and whole genome libraries; K.L. and H.H. discussed results and provided subject-matter expertise; J.S. and S.C. conceived the overall study and were responsible for the final version of the manuscript. All authors read and approved the final manuscript.

## Figure Legends

**FIG S1** Relative abundances of 4 major bacterial phyla across 14 skin sites from normal human volunteers. Relative abundances determined by sequencing the V1-V3 region of 16S rRNA followed by classification with the DADA2 and the eHOMD v15.1 database.

**FIG S2** Relative abundance of the major *C. tuberculostearicum* ASVs across 14 skin sites in normal human volunteers.

**FIG S3** A schematic representation of 16S rRNA intra-genome variation. Gray rectangles represent the complete V1-V3 region of each of the four 16S rRNA gene copies found in each genome. Variant positions are marked with an X and colored blue for variants characteristic of ribotype B, or orange for additional variants or variants that are found outside the trimmed ASV (red dotted line).

**FIG S4** Alignment of the *C. tuberculostearicum* reference genome with five PacBio genomes from this study. Aligned regions are shown as bands colored by the percent identity.

**FIG S5** Average Nucleotide Identity (ANI) of *C. tuberculostearicum* species complex genomes. The distance matrix was calculated using fastANI and bidirectional percent identities were averaged. Distances were hierarchically clustered and visualized using heatmap.2/R. Species are abbreviated as *C. tuberculostearicum*, Ctub; *C. curieae*, Ccur; *C. kefirresidentii*, Ckef. Pairwise comparisons at >95% identity are marked with a (*).

**Fig S6** Functional classifications of orthologous gene clusters. Dotted line demarcates expected proportion of core, accessory, and singleton genes assigned to each category given the total (N=3703 including 243 duplicate and triplicate assignments) and core (N=1738, or 47%) number of category assignments. Total gene counts per category *N* are labeled.

**TABLE S1** Table of dereplicated whole genomes. N50 is the N50 contig length, n/a for finished genomes. The number of plasmids are listed for finished genomes. Except for CTNIH9 (*), all genomes are from strains associated with HVs in the microbiome analysis. ASV, Amplicon Sequence Variant; HV, Healthy Volunteer

**TABLE S2** Phage defense systems and methylation patterns for complete *Corynebacterium tuberculostearicum* species complex genomes. Defense finder reports restrictions modification systems (RM) and other phage defense systems. Numbers in parenthesis indicate the number of systems present, if more than one.

**TABLE S3** Ribotype-specific genes. Genes uniquely present among all ribotype A or ribotype B genomes. All ribotype:gene associations p-values (FDR-adjusted) < 0.05.

